# TEgenomeSimulator: A Flexible Framework for Simulating Genomes with Configurable Transposable Element Landscapes

**DOI:** 10.64898/2026.03.09.710711

**Authors:** Ting-Hsuan Chen, Olivia Angelin-Bonnet, James Bristow, Christopher Benson, Shujun Ou, Cecilia Deng, Susan Thomson

## Abstract

Transposable elements (TEs) are major contributors to genome structure and evolution. However, our ability to study them is limited by the difficulty of annotating and curating them, especially in non-model organisms. This challenge is compounded by a lack of ground-truth datasets for benchmarking, which are nearly impossible to generate through manual curation alone. To overcome this critical limitation, we developed TEgenomeSimulator, a flexible framework for generating synthetic genomes with configurable TE landscapes. TEgenomeSimulator supports both randomly generated and biologically derived backbone sequences, enabling the modeling of TE insertions under diverse structural and evolutionary contexts. Benchmarking against existing simulators demonstrated that TEgenomeSimulator reproduces realistic TE composition, sequence divergence, and integrity distributions while offering greater flexibility in modeling chromosome-level structure. Its modular design offers a tuneable continuum between biological fidelity and experimental control, facilitating systematic benchmarking, algorithm development, and evolutionary modeling of TE dynamics *in silico*, filling a major gap in the field. The source code of TEgenomeSimulator and the scripts for this paper are available at https://github.com/Plant-Food-Research-Open/TEgenomeSimulator.

## Introduction

Transposable elements (TEs) are abundant components in eukaryotic genomes. Once regarded as parasitic DNA, TEs are now recognized as major contributors to gene regulation, genome architecture, and evolutionary processes [1,2]. TEs also impact agronomic traits, such as plant-pathogen interactions and environmental responses [3], and may act as “silent” regulators of chromosomal architecture and post-zygotic barriers that drive speciation [4]. Consistent with this view, cross-taxon surveys demonstrate that genome size variation can be largely explained by TE accumulation [5-9]. TEs have also been implicated in centromere structure evolution [6,7] and higher-order chromatin architecture [12]. Together, these discoveries establish TEs not only as regulators of genes but also as major catalysts of genome evolution, underscoring the need for high-quality assemblies to accurately resolve repetitive DNA.

Advances in long-read sequencing technologies and assembly algorithms have markedly enhanced genome assemblies and the resolution of repetitive regions, leading to increased genome and pan-genome assemblies across taxa. Recent pangenome studies in *Brassica*, grapevine and potato highlight TE enrichment in dispensable genomic regions linked to resistance gene family expansion, regulatory variation, and reproductive isolation [13–15], further solidifying the structural and functional impact of TEs and their critical role in genome evolution. Despite these advances, TE annotation and interpretation remain underprioritized in many genome projects, in only a few cases have domain experts filled this knowledge gap [11].

The detection and annotation of TEs within assembled genomes has benefited from bioinformatic tool developments [12]; however, there are still major challenges to generating quality annotation, especially for non-model organisms. TEs vary greatly in repetitiveness and have undergone mutation, nested TE insertion and ectopic recombination, complicating the differentiation of aged TE copies from non-TE sequences and hindering accurate reconstruction of the consensus TE sequences [12]. Besides, ground–truth datasets are especially scarce for non-model organisms, and only a small number have had any validation through other ‘omics’ data sources [13,14]. Dfam is a TE database with an open framework for community contribution and transparency in curated and non-curated datasets [16]. As of March 2025, only 0.6% of the TE families in Dfam (release 3.9) have been manually curated, predominantly from mammals. Most curated libraries rely on labor-intensive manual curation [17]. Consequently, simulated genomes with traceable TE insertions can address this bottleneck by providing benchmarks for evaluating TE detection and annotation.

Simulating TE sequence at scale that reflects the breadth of complexity and diversity of TE composition is essential for benchmarking TE detection tools and modeling TE activity over evolutionary time. Synthetic genomes with deliberately designed TE architectures, including nested insertions, target site duplications (TSDs) and extreme cases of sequence divergence and polymorphisms, provide a controlled framework to probe the limits of TE detection and classification methods. This is particularly crucial in the era of telomere-to-telomere assemblies and pangenome-based annotations. By generating such comprehensive and customizable datasets, researchers can systematically evaluate performance under realistic and challenging conditions while also exploring the evolutionary processes shaping genome architecture. A variety of TE simulators have been developed to address some of these needs. For example, denovoTE-eval [18] synthesizes random sequences with user-specified lengths and GC content, requiring an input table containing TE families and parameters to model mutagenesis, sequence decay, and random insertions. However, it can simulate only a single random sequence at a time and offers limited flexibility for adjusting mutagenesis parameters when working with large TE libraries. By relying on predefined TE architectures, SimulaTE [19] can recapitulate intricate TE landscapes and simulate sequencing reads, but it remains limited in its capability to simulate novel evolutionary events that extend beyond the user-provided genome architecture. SLiM [20] excels at modeling selection and demography but represents TEs as single units, each like a single nucleotide variant, restricting exploration of sequence-level architectures and genome size changes. GARLIC [21] generates realistic intergenic sequences based on k-mer composition, GC content, and repeat profiles but lacks flexibility in TE mutagenesis profiles and does not capture the coexistence of coding and non-coding regions. While PrinTE is designed for forward-time simulation on TE mobilization, mutation, and removal, it requires multiple iterative attempts to infer and reproduce the dynamics of a specific TE type, with a focus on LTR retrotransposon [22]. Despite the breadth of existing simulation tools in design, few offer the flexibility to customize genome content while also modeling TE sequence mutagenesis under diverse decay schemes. Even fewer support fine-grained, family-specific controls or allow combinations of multiple modes for complex TE composition simulations.

To address these limitations, we developed TEgenomeSimulator (**Fig. 1**), which was built on concepts introduced in denovoTE-eval. Extending beyond the capability of denovoTE-eval in synthesizing single random sequences and performing random and nested TE insertions, TEgenomeSimulator can synthesize genomes with multiple chromosomes of varying complexity and offers fine-grained control over TE sequence populations for a more realistic fragmentation pattern. TEgenomeSimulator departs from the uniform TSD range approach of denovo-TEeval by modeling TSD length at the TE-superfamily level, a key feature for structure-based TE detection and classification. It can also incorporate real genome assemblies and their TE compositions for more realistic modeling. Furthermore, it allows users to combine different simulation modes to introduce new TE activity beyond known TE compositions, enabling the exploration of diverse evolutionary scenarios. A comparison with existing simulators is available in **Table S1**.

**Figure 1.**
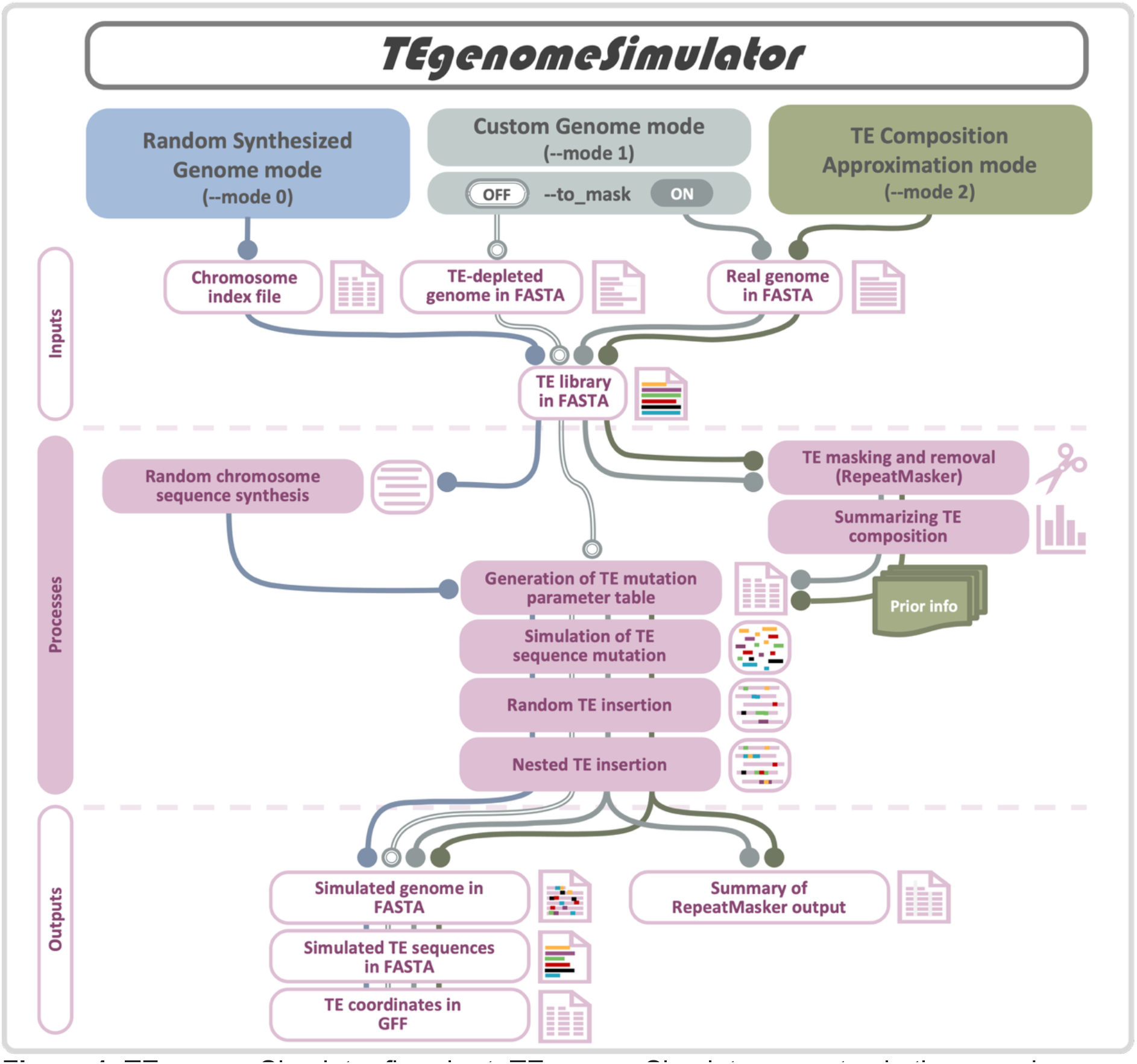
TEgenomeSimulator flowchart. TEgenomeSimulator operates in three modes— **Random Synthesized Genome** (**mode 0**), **Custom Genome** (**mode 1**), and **TE Composition Approximation** (**mode 2**)—shown in blue, grey, and sage, respectively. Workflow components (rounded pink rectangles) are organized into three conceptual layers: Inputs, Processes, and Outputs. Connections between components indicate the data flow and processing steps taken by each mode, highlighting shared and mode-specific elements. See Materials and Methods for details.

In this paper, we demonstrate that TEgenomeSimulator generates more realistic synthetic genomes than denovoTE-eval and GARLIC. We systematically illustrate how parameterization of TE copy number, fragmentation, mean sequence identity, and standard deviation (SD) influences simulation outcomes and their biological interpretation. The resulting outputs enable robust benchmarking of TE detection and annotation methods. As a proof of concept, we use TEgenomeSimulator to facilitate the evaluation of TE detection by RepeatMasker [23]. TEgenomeSimulator is implemented as a reproducible and portable Python package for Linux, installable via ‘pip’, Docker, or Apptainer.

## Methods

### Implementation

#### Simulation Modes

Random Synthesized Genome mode (**mode 0**) synthesizes artificial chromosomes using user-defined chromosome lengths and GC content, then inserts simulated TEs at random positions. A TE library is used to produce TE copies subjected to random mutagenesis, including nucleotide substitutions, single-base insertions and deletions (InDels), TSDs, and fragmentation.

Custom Genome mode (**mode 1**) accepts a user-provided ‘backbone’ genome with TEs removed, or masks and removes TEs using RepeatMasker [23] (using ‘--to_mask’ option) with a user-supplied TE library. Mutation and insertion proceed as in **mode 0**.

TE Composition Approximation mode (**mode 2**) also utilizes RepeatMasker to remove TEs but additionally infers TE family abundance, nucleotide substitution and InDel rates, and sequence integrity from the source genome, which are then used to parameterize TE simulations.

#### Simulating TE Mutagenesis

From the TE library, TEgenomeSimulator extracts family classification and sequence metadata and assigns per-family mutagenesis parameters. In **modes 0** and **1**, per-family TE copy numbers are sampled from user-defined ranges. Sequence identity (*I*) is drawn from a Gaussian distribution with user-specified mean and SD (**modes 0** and **1**) or estimated from RepeatMasker output (**mode 2**), then adjusted following denovoTE-eval. TSD lengths follows published observations [24], with random TSD sequences synthesized. Sequence integrity in **modes 0** and **1** is modeled using a beta distribution controlled by alpha and beta (default α=0.5, β=0.7), producing predominantly fragmented elements, with 0.1% of copies remaining intact by default. Sequence integrity in **mode 2** is inferred from the source genome. TE copies are randomly oriented and integrated. See **Supplementary Methods** for detailed implementation.

#### Random and Nested TE Insertion

Nested insertions are generated in 0-30% of LTR retrotransposon copies, with insertion targets and locations selected randomly.

#### Input and Output

TE libraries must follow RepeatMasker naming conventions (e.g, >ATCOPIA10#LTR/Copia) and ideally contain full-length sequences (e.g. LTR-INT-LTR for retrotransposons). Split LTR entries are automatically reassembled (see **Supplementary Methods**). **Modes 1** and **2** require a genome FASTA file, whereas **mode 0** requires a genome index CSV file specifying chromosome name, length, and GC%. Outputs include the simulated genome, all TE copy sequences, and a GFF file recording TE locations, identity, integrity, and nesting relationships.

### Simulator comparison analysis

#### Comparison 1: TEgenomeSimulator vs denovoTE-eval

TEgenomeSimulator’s **mode 0** was used to synthesize a 10 Mb non-TE random sequence with 35% GC content, followed by TE insertion using a composition defined by TEgenomeSimulator (denoted as TEGS in **Fig. 2**). Copy numbers per family were set between 1 and 10, with other mutagenesis parameters left at default values. To ensure comparability, settings for size of non-TE backbone, GC%, TE copy number, and sequence identity from TEgenomeSimulator were used for denovoTE-eval (denoted as dnvTEe in **Fig. 2**), while maintaining direct comparisons of other features.

**Figure 2.**
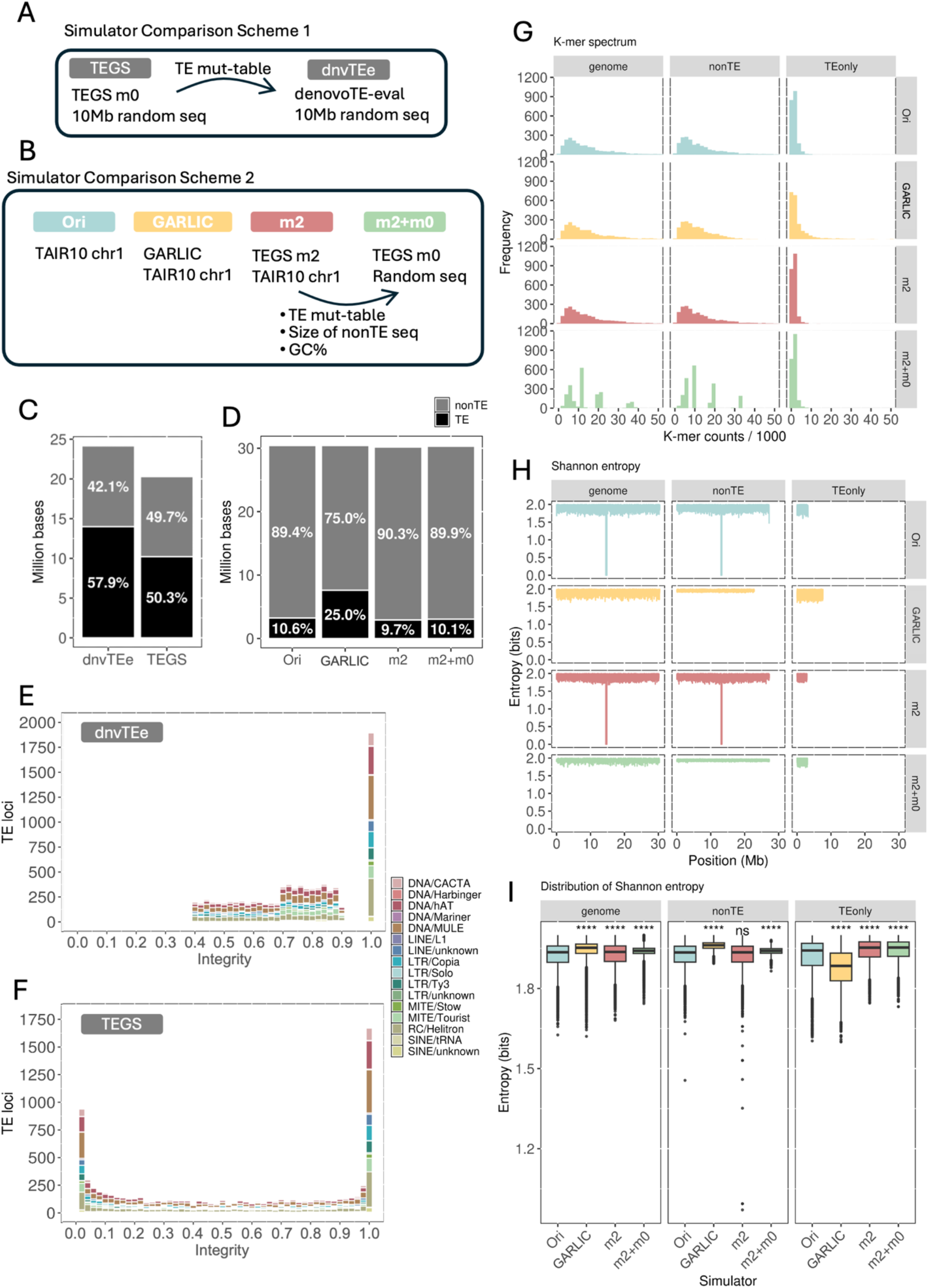
Simulator Comparisons. (**A**) Comparison 1: **mode 0** of TEgenomeSimulator (TEGS) and denovoTE-eval (dnvTEe) were compared using a random synthesized 10 Mb sequence. The TE mutagenesis table (TE mut-table) generated by TEGS was imported to dnvTEe to match TE composition for comparison. (**B**) Comparison 2: simulated TAIR10 chromosome 1 (chr1) generated by GARLIC, TEGS full **mode 2** (m2), TEGS partial **mode 2** followed by full **mode 0** (m2+m0), and the original TAIR10 chr1. For m2+m0, the TE muttable, TE-depleted chr1 length and GC content were acquired from m2 and used as inputs for m0. (**C, D**) Proportions of TE and non-TE sequences from Comparison 1 (C) and 2 (D). (**E, F**) TE integrity distributions for dnvTEe (E) and TEGS (F) simulations. (**G, H**) K-mer (G) and Shannon entropy (H) spectra of whole genomes, and the non-TE and TE only sequences, colored by simulation method. (**I**) Shannon entropy ranges by sequence category. Asterisks indicate *p* < 0.0001 (Wilcoxon Rank-Sum Test versus the original genome).

#### Comparison 2: TEgenomeSimulator vs GARLIC

Chromosome 1 of the *Arabidopsis thaliana* (TAIR10, NCBI accession ID GCF_000001735.4) and the curated TE library from the EDTA GitHub repository (https://github.com/oushujun/EDTA/blob/master/database/) were used as inputs. Two genomes were simulated with TEgenomeSimulator: (i) a digital replica of TAIR10 chromosome 1 using **mode 2** (m2 in **Fig. 2**); and (ii) a hybrid simulation in which the TE composition inferred by **mode 2** was imported into **mode 0** to generate a randomly synthesized backbone (m2+m0 in **Fig. 2**; see **Supplementary Methods**).

GARLIC simulation required four input files: (i) tandem repeat profiles generated by Tandem Repeat Finder (TRF, v4.07) [25] on TAIR10 genome; (ii) gene annotations from Ensembl release 61 (GFF3) converted to GARLIC’s tabular format using a custom script; (iii) RepeatMasker’s alignments of TAIR10 genome using the same TE library; and (iv) the TE library converted from FASTA to EMBL format. GARLIC’s ‘createModel.pl’ and ‘createFakeSequence.pl’ scripts were used to generate a synthetic sequence matching chromosome 1 length.

K-mer spectra (k = 6) were analyzed using a custom Python script treating reverse complements as canonical. Shannon entropy (*H*) on nucleotide probabilities (*p*) was calculated in 1 Kb sliding window with a step size of 1 bp according to the formula:

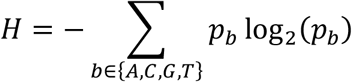

### Use-cases demonstration

A multi-species TE library was constructed by merging the curated TE library of *A. thaliana, Oryza sativa*, and *Zea mays* obtained from EDTA, followed by redundancy removal using cd-hit-est with settings following Goubert et al. [17]. The resulting library, along with the TE-depleted TAIR10 genome, was used for simulation demonstrations using **mode 1**. The TAIR10 genome was pre-processed using RepeatMasker and the curated *A. thaliana* TE library to remove endogenous TEs. Simulation settings corresponding to **Fig. 3** are described below.

**Figure 3.**
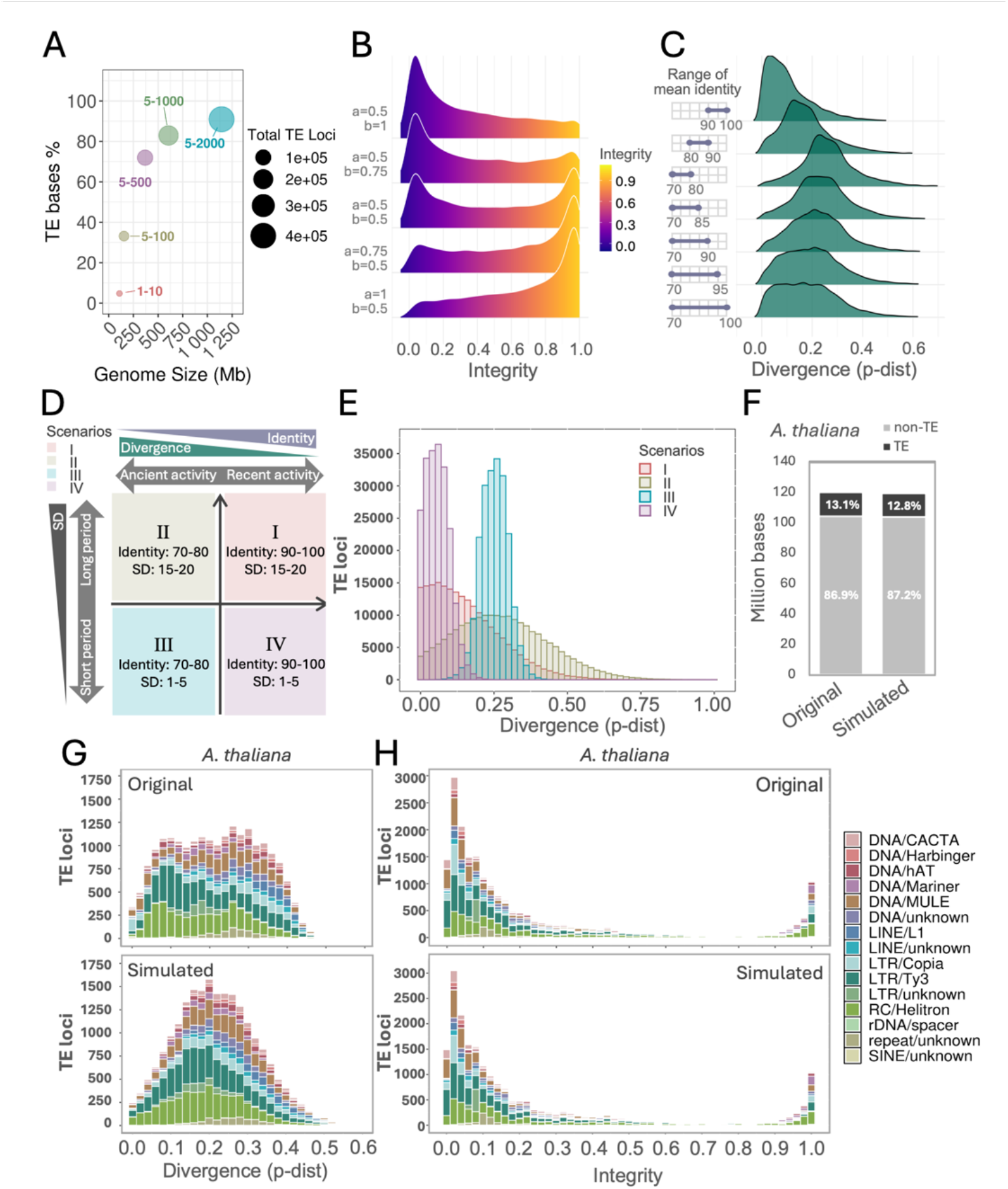
Example use-case of TEgenomeSimulator. (**A**) Impact of altering TE copy number ranges on simulated genome size (**mode 0**). (**B**) TE sequence integrity distribution under different alpha and beta settings (**mode 0**). (**C**) TE sequence divergence distribution under varying ranges of mean identity across TE families, with a fixed TE copy number range of 5–100 (**mode 0**). (**D**) Four historical TE activity scenarios modeled by adjusting mean identity (activity timing) and its standard deviation (SD; activity duration). (**E**) Resulting TE divergence patterns associated with the settings in (C). (**F**) TE/nonTE sequence proportions in original and simulated *Arabidopsis thaliana* genomes (**mode 2**). (**G**,**H**) Sequence divergence (G) and integrity (H) in original and simulated *A. thaliana* genomes.

#### Impact of TE copy number on genome size (mode 1)

TE copy number ranges per family were set to 1-10, 5-100, 5-500, 5-1000 and 5-2000 using the TE-depleted *A. thaliana* genome as input.

#### Impact of alpha and beta on TE integrity (mode 1)

For demonstration, TE integrity was modeled with a beta distribution controlled by five [α, β] pairs: [0.5, 1], [0.5, 0.75], [0.5, 0.5], [0.75, 0.5], and [1, 0.5].

#### Impact of mean sequence identity (mode 1)

Identity ranges were set to 90-100, 80-90, 70-80, 70-85, 70-90, 70-95, and 70-100.

#### Combined effects of sequence identity and standard deviation (mode 1)

Four scenarios were simulated (**Fig. 3D**) by varying mean identity ranges (10-80 or 90-100) and SD (1-5 and 15-20) to represent different histories of TE activity (see **Supplementary Methods**)

#### Digital replicas (mode 2)

Genomes of *A. thaliana* (TAIR10; 120 Mb), *O. sativa* (AGIS1.0; NCBI accession ID GCF_034140825.1; 386 Mb), *Z. mays* (Zm-B73-REFERENCE-NAM-5.0; NCBI accession ID GCF_902167145.1; 2.183 Gb), *Danio rerio* (GRCz12tu; NCBI accession ID GCF_049306965.1; 1.449 Gb) and *Drosophila melanogaster* (dm6; NCBI accession ID GCF_000001215.4; 138 Mb) were directly used as input with default settings. Acquisition of TE libraries for the mentioned plant species has been described previously. TE libraries for *D. rerio* and *D. melanogaster* were downloaded from the RepeatModeler2 GitHub repository (https://github.com/jmf422/TE_annotation/tree/master/benchmark_libraries/RM2).

### Computational resource tests

**Mode 0** was benchmarked using five configurations that varied the length of randomly synthesized sequences with 35% GC prior to TE insertion. Four tests, generated a single sequence of 100 Kb, 1 Mb, 10 Mb, or 100 Mb, and a fifth generated ten 100 Mb sequences (1Gb in total). TE copy numbers per family were fixed at 1-10.

To test **mode 1**, the TE-depleted TAIR10 genome and curated *A. thaliana* TE library were used, with TE copy number ranges of 1-10, 5-100, 5-500, 5-1000, and 5-2000.

**Mode 2** resource usage was recorded for simulations of *A. thaliana, O. sativa, Z. mays, D. melanogaster*, and *D. rerio*.

All tests were executed on the same SLURM-managed cluster. Each **mode 0** and **mode 1** test used a single CPU core. Each **mode 2** test was assigned 20 CPU cores due to the computational demands of running RepeatMasker.

### TE recovery rate analysis

RepeatMasker was applied to simulated genomes from the four scenarios **Fig. 3D** using the same TE library as in the simulations. TEgenomeSimulator’s TE annotation files (GFF) served as ground truth and were compared to the BED files converted from RepeatMasker output using Bedtools Intersect [26], applying an 80% reciprocal overlap cutoff. Recovery rate (*R*) was calculated per integrity and identity bin (10 bins, ranging from 0 to 1) according to the formula:

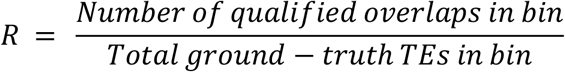

## Results

### Simulator Comparisons

We compared the key features of TEgenomeSimulator to those of denovoTE-eval, GARLIC, SimulaTE, and SLiM (**Table S1)**. Unlike denovoTE-eval, TEgenomeSimulator enables the synthesis of multi-chromosomal genomes within a single simulation run. It also supports modeling of superfamily-specific TSD lengths, allows beta distribution or empirical distribution to model fragmentation, and enables both empirical and combinatorial modeling of TE composition through the sequential application of multiple simulation modes (**Supplementary Methods**). In contrast to GARLIC, which analyzes genic and non-genic profiles but only synthesizes non-genic regions, TEgenomeSimulator’s **mode 2** preserves gene models whose exons or introns are not identified and masked as TEs by RepeatMasker. Moreover, TEgenomeSimulator is uniquely capable of utilizing either randomly generated or empirically derived non-TE backbones for TE insertion, providing enhanced flexibility for the simulation of complex and biologically realistic TE landscapes (**Table S1**).

We assessed TEgenomeSimulator against its closest peers, denovoTE-eval and GARLIC, using two comparison designs (**Fig. 2A-B**). Comparison 1 (**Fig. 2A**) utilizes a 10 Mb TE-free backbone, synthesized by TEgenomeSimulator, that is used by both TEgenomeSimulaor and denovoTE-eval for subsequent TE insertion (see **Methods**). To maintain consistency, the TE mutagenesis parameters of TEgenomeSimulator were reformatted and imported into denovoTE-eval. Both tools yielded comparable TE superfamily composition and divergence distributions (**Fig. S1**), validating the consistency in the transferred TE composition and sequence identity parameters. We did, however, observed considerable variance in the resulting genome size and composition. denovoTE-eval produced a genome approximately 5 Mb larger than TEgenomeSimulator, due to incorporating 7.6% more TE bases(**Fig. 2C**). We also observed a marked difference in the integrity distributions of simulated TEs. denovoTE-eval displayed flattened TE loci distributions at integrity ranges 0.4-0.7 and 0.7-0.9, with no fragmented element exceeding 0.9 (**Fig. 2E**). This gap between 0.9 and 1 arose because denovoTE-eval caps integrity at 0.9 following fragmentation, while the even patterns reflected its programmed two-tier sampling scheme: integrities 0.7-0.9 for sequence <500 bp and 0.4-0.9 for sequences ≥500 bp. TEgenomeSimulator, however, models integrity using a beta-distribution combined with a fixed fraction of intact TEs. This results in a broader spectrum, spanning degraded to nearly intact elements, better reflecting the sequence integrity observed in natural genomes (**Fig. 2F**).

For Comparison 2 (**Fig. 2B**), we used TEgenomeSimulator’s **mode 2** to build a model of the TE content of TAIR10 chromsome1 (chr1), which was then used as input for TE insertion into TE-depleted chr1 (m2) or a simulated backbone matching the length of chr1 (m2+m0). The resulting genomes were compared to simulations derived from GARLIC. GARLIC produced genomes with a markedly higher proportion of TE sequence (25%) than the original chr1 (10.6%) and nearly three times as many TE insertions (**Fig. S2A**). Both the m2 (9.7%) and m2+m0 (10.1%) genomes more closely matched the balance of TE and non-TE occupancy observed in chr1 (**Fig. 2D**). GARLIC’s integrity profile was heavily skewed toward values of 0.9-1, in sharp contrast to the more realistic integrity ranges captured by TEgenomeSimulator m2 and m2+m0 (**Fig. S3**). Sequence complexity analyses reinforced these differences (**Fig. 2G-I**). GARLIC’s k-mer profile displayed an extended tail in TE-only regions, which is absent from both chr1 (Ori) and TEgenomeSimulator-synthesized sequence (**Fig. 2G**). In non-TE regions, GARLIC and TEgenomeSimulator’s m2 closely matched the reference k-mer profile, whereas the distortion in m2+m0 reflected the use of a randomly synthesized non-TE backbone (**Fig. 2G**). We used Shannon entropy analysis to determine variability within the data, we observed a central dip within chr1 (Ori) that was successfully reproduced by TEgenomeSimulator’s m2, while GARLIC and m2+m0 failed to capture this feature (**Fig. 2H**). GARLIC’s outputs exhibited a narrower, higher entropy range in non-TE regions and a lower entropy range in TE-only regions relative to the original genome (**Fig. 2I**). For TEgenomeSimulator, the TE-only entropy range of m2 was shifted compared to the reference, but its non-TE profile was preserved, while m2+m0 showed markedly narrower non-TE distribution, again reflecting the random backbone synthesis (**Fig. 2I**).

### Use cases

TE composition is shaped by multiple evolutionary processes, each leaving distinct genomic signatures. To illustrate how TEgenomeSimulator captures these dynamics, we demonstrate key configurations that reproduce major features of empirical TE landscapes.

#### Modeling basic TE features

TE accumulation is a primary driver of genome expansion [27–29]. Using the Custom Genome Mode together with the TE-depleted *A. thaliana* (TAIR10) genome and the curated *Arabidopsis* TE library (see **Methods**), we simulated genome expansions ranging from under 200 Mb to over 1 Gb by increasing the maximum copy number per TE family from 10 to 2000 (**Fig. 3A**). This configuration directly models the impact of TE accumulation on genome size.

TE integrity reflects the balance between recent insertions and long-term decay. In both Random Synthesized Genome and Custom Genome modes, TE integrity is modelled using a beta distribution across families. When α < β (both between 0 and 1), the distribution takes on an asymmetrical U-shape with a peak near low integrity values (**Fig. 3B**), reflecting the predominance of TE relics observed in empirical datasets [30]. Conversely, when β < α, the distribution shifts toward high-integrity copies (**Fig. 3B**), reflecting recent evolutionary TE activity.

Alongside modifying TE integrity, we can vary the range of mean sequence identity to reflect mutation accumulation over time. By varying the range of mean sequence identity (70%-100%) while constraining copy numbers (5-100 per family), we generated divergence distributions with progressively flattened peaks as identity ranges widened (**Fig. 3C**), reproducing the expected signatures of sustained or overlapping TE amplification events [31,32].

### Modeling TE composition under diverse evolutionary scenarios

The duration of TE bursts, which can be measured by the standard deviation (SD) of sequence identity, also influences divergence distribution. A short-lived TE burst results in a rapid deposition of nearly identical insertions. These insertions then gradually diverge through accumulated mutation. In contrast, a long-lasting burst over evolutionary time leads to the continuous deposition of similar TE sequences, causing a less synchronized divergence pattern owing to overlapping degradation timelines. Moreover, ancient TE bursts tend to show greater sequence deterioration due to prolonged exposure to mutational decay. We can simulate genomes that represent different evolutionary phases of TE activity (**Fig. 3D**), offering genomic templates with distinct base-level TE composition that can serve as burn-in states for more advanced simulations. As shown in **Fig. 3D**, the representation of TE burst timeline and duration on the x-axis and y-axis, respectively, allows four distinct TE divergence landscapes to emerge: I) high sequence identity with high SD, modeling recent TE bursts with ongoing or prolonged activity over time; II) low sequence identity with high SD, modeling ancient and prolonged activity, or multiple independent bursts occurring over an extended period; III) low sequence identity with low SD, reflecting ancient TE bursts that were quickly inactivated, with TE activity remaining silenced since the initial event; IV) high sequence identity with low SD, inferring very recent and rapid TE bursts, resulting in a synchronized insertion of highly similar sequences.

We modelled these four scenarios using the Custom Genome mode of TEgenomeSimulator. The identity and SD ranges were implemented as shown in **Fig. 3D**, with identity ranges of 90%–100% and 70%–80%, and SD ranges of 15–20 and 1–5. The TE copy number range was fixed between 5 and 1000. The resulting sequence divergence profiles demonstrated a narrowing of the peak as the SD range decreased from 15–20 to 1–5, indicating more instantaneous bursts (**Fig. 3E**). Furthermore, shifting the identity range from 70–80% to 90– 100% led to a leftward shift of the divergence peak (from ~0.25 to ~0.1) and an increase in peak height. These results demonstrate that TEgenomeSimulator is well-suited for modeling TE composition under diverse evolutionary scenarios.

#### Digital replicas

TE composition in real genomes varies substantially across TE families and species because of their complex and diverse evolutionary histories. This variability makes it challenging to simulate realistic genome-wide TE landscapes. To address this, we used TEgenomeSimulator’s TE Composition Approximation Mode to model the TE profiles of *A. thaliana* (TAIR10), *O. sativa, Z. mays, D. rerio*, and *D. melanogaster* and synthesis replicas. We observed the final genome size and proportion of TE occupancy in the digital replicas closely matched those observed in the original genomes (**Fig. 3F, Fig. S4-S7**). In terms of TE loci and TE bases per superfamily, the simulated TE composition also aligned well with the original genomes (**Fig. S4-S8**).

While the simulated divergence profiles of the replicas resembled those of the original genomes, they were not identical: the fine-scale fluctuations present in the original genomes were smoothed in the replicas due to the use of normal distributions in the simulator. This approach, however, still provides a generalized distribution pattern that captures the major trends in TE sequence evolution (**Fig. 3G, Fig. S4-S7**).

To replicate a real TE integrity pattern for each TE family, the empirical distribution of TE length ratios (relative to consensus sequences) is extracted from the input genome. This enables the simulator to reproduce the integrity distributions more authentically in the resulting simulated genome (**Fig. 3H, Fig. S4-S7**).

These digital replicas provide a robust foundation for simulating future TE-related events, such as new TE bloating or purging episodes, effectively serving as burn-in phases for modeling genome evolution from the current state.

### Computational resource usage

To evaluate the computational resource usage of TEgenomeSimulator, we conducted tests across all three simulation modes (**Fig. 4, Table S2**). In **mode 0**, we observed that the total length of randomly synthesized sequences significantly impacts memory usage, particularly for sequences longer than 100 Mb. Synthesizing a large genome over 1 Gb, comprising 10 chromosomes of approximately 100 Mb each, required over 12 GB of memory (**Fig. 4, Table S2**). In **mode 1**, increasing the TE copy number range per family had a more pronounced impact on runtime than on memory usage (**Fig. 4**). Simulations with broader copy number ranges took several hours to complete and required multiple gigabytes of memory, depending on the settings. Notably, a simulation with copy number ranges of 5-2000 per family consumed over 6 GB of memory and required nearly 10 hours to complete, resulting in a simulated genome exceeding 1 Gb in size (**Fig. 4, Table S2**). In contrast, tests varying TE sequence identity and integrity finished in under 3.5 minutes and used less than 1 GB of memory (**Table S2**). In **mode 2**, both runtime and memory usage were substantially increased in relation to the size and complexity of the input genomes used for TE composition approximation (**Fig. 4, Table S2**). The increasing computational demand with genome size appears to correlate with native TE content (**Fig. S2-S5**).

**Figure 4.**
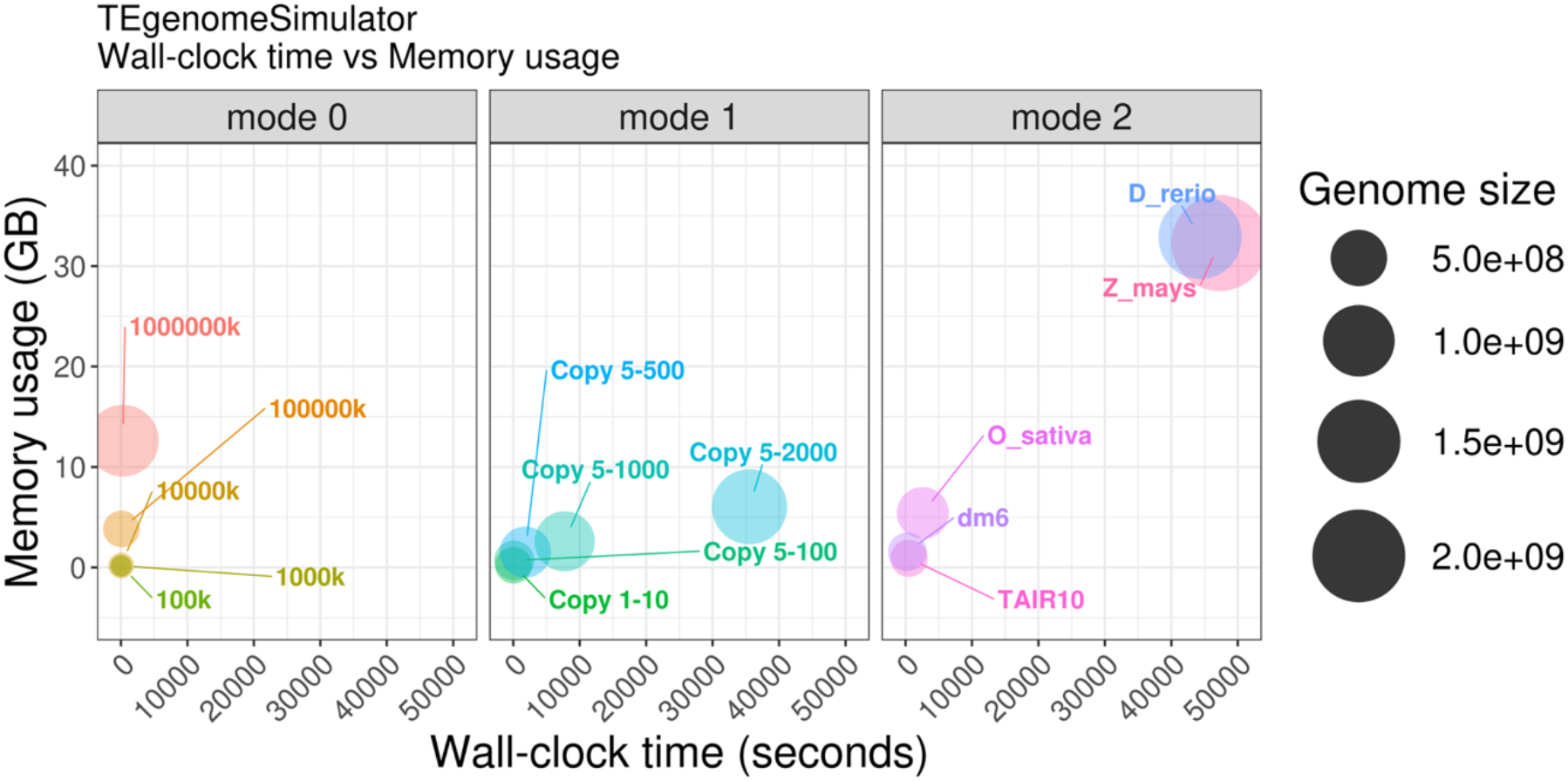
Computational resource usage of TEgenomeSimulator. Each bubble represents a test run, with size proportional to the simulated genome. **Mode 0** varied reference sequence length (100 Kb-100 Mb). **Mode 1** varied TE copy numbers ranges. **Mode 2** approximated TE compositions from five genomes: *Arabidopsis thaliana* (TAIR10), *Oryza sativa* (O_sativa), *Zea mays* (Z_mays), *Drosophila melanogaster* (dm6), and *Danio rerio* (D_rerio).

### TE recovery rate test

TEgenomeSimulator can generate diverse TE insertions ideal for benchmarking TE detection and annotation tools. We assessed the TE detection capability of RepeatMasker using simulated TE insertions from the four scenarios described in **Fig. 3D** as ground truth (**Fig. 5A**). TE recovery rate was measured as the proportion of simulated insertions correctly identified by RepeatMasker, stratified by sequence identity and integrity (**Fig. 5A**). The resulting heatmaps revealed a strong dependence on sequence identity, with higher recovery for insertions of greater similarity to the consensus (**Fig. 5B**). In contrast, recovery showed little or no clear dependency on integrity. Bar plots summarizing recovery rate across identity and integrity slices confirmed these trends across the four scenarios: recovery rates remained high at identity above 0.7 but declined progressively with greater divergence, whereas variation across integrity levels lacked a consistent pattern. Although the precise recovery values may be influenced by RepeatMasker’s parameter settings, particularly the divergence and alignment thresholds, the results clearly demonstrate that TE insertions with higher sequence divergence are more difficult to detect.

**Fig. 5.**
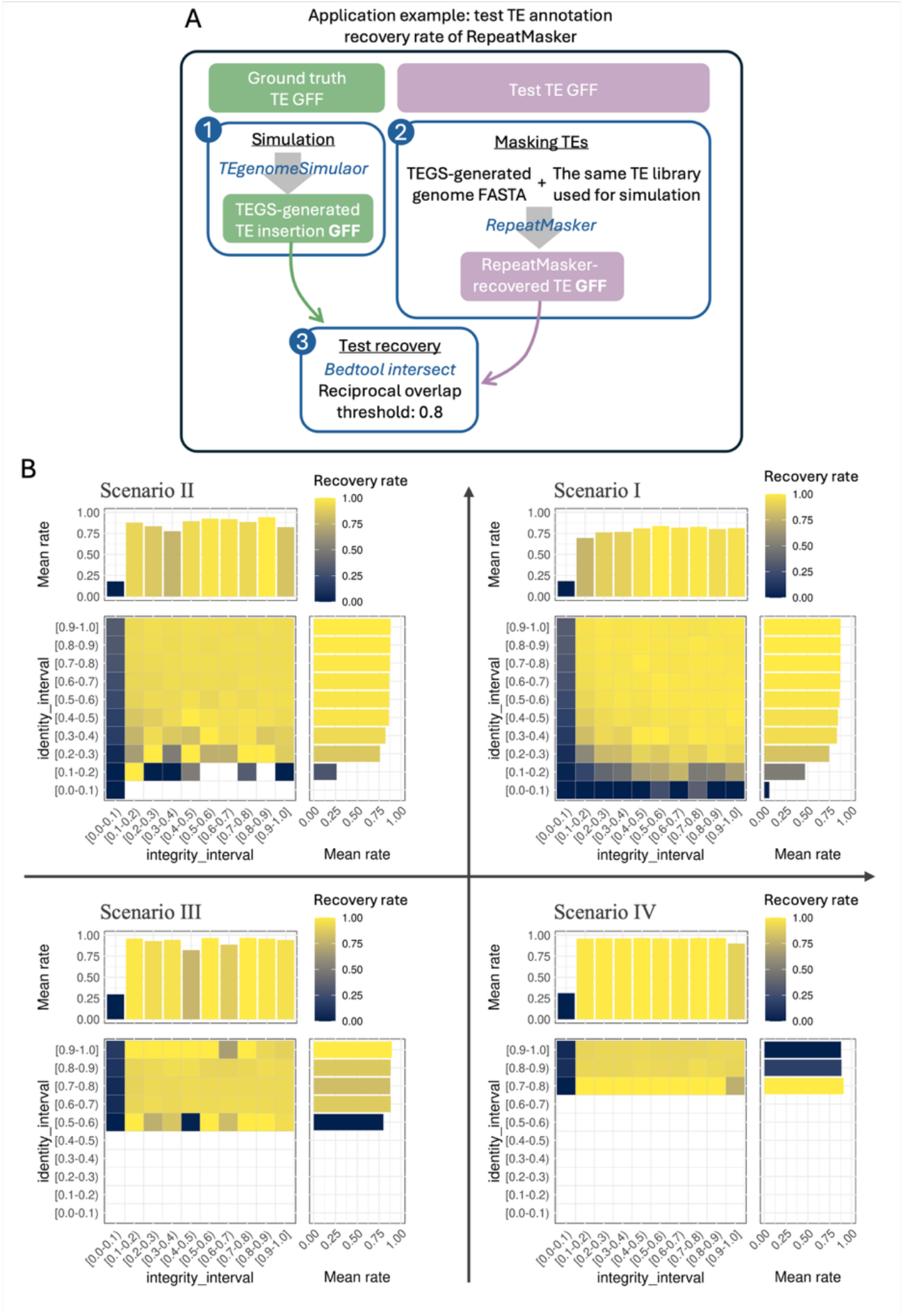
TE annotation recovery rate of RepeatMasker. (**A**) TE annotation recovery workflow, with ground-truth (green) and test (purple) annotations; numbered blue rectangles indicate analysis steps (see **Methods**). (**B**) Heatmaps of recovery rate by sequence identity and integrity for the four scenarios in **Fig. 3D**, with marginal bar graphs showing mean recovery. Empty bins are shown in white.

## Discussion

TEgenomeSimulator introduces a flexible, mode-based architecture that addresses critical limitations in existing TE simulation tools by enabling both randomized simulations and simulations based on real genomic architecture, thereby capturing structural and compositional complexity characteristics of natural TE landscapes.

TEgenomeSimulator-generated TE populations display a broad and continuous range of sequence integrity values by modeling either a beta or an empirical distribution, providing a more biologically realistic material for testing annotation tools. While denovoTE-eval served as a foundational model for the development of TEgenomeSimulator, it lacked flexibility in defining superfamily-specific TSDs and exhibited discontinuous integrity distributions due to rigid random sampling. TEgenomeSimulator overcomes this limitation by introducing TE superfamily modeling of TSDs and sequence divergence and integrity, producing outputs that better reflect natural TE diversity and evolutionary behaviors of TE lineages observed in real genomes. By comparison, GARLIC’s emphasis on modeling non-genic contexts is useful for certain studies but results in overrepresentation of high-integrity TE copies and reduced sequence entropy, limiting its value for representing complex genomic structure and simulating evolutionary variation. Other simulation tools, such as SimulaTE and SLiM, emphasize population dynamics or evolutionary events but lack fine control over structural variation or sequence-level mutagenesis parameters that TEgenomeSimulator provides.

TEgenomeSimulator’s capability in combining different modes provides a tunable continuum across a spectrum of evolutionary scenarios and genomic realism. For example, a **mode 0** simulation is ideal for controlled benchmarking under a uniform backbone condition, where the goal is to assess TE detection sensitivity without confounding genomic context effects. **Mode 2**, approximation to realistic TE composition, best supports validation of annotation tools with authentic genomic contexts, where chromosomal architecture, including centromere, intergenic and intragenic regions, is preserved as the non-TE backbone. **Mode 1** combines controllable TE composition and mutagenesis with a realistic non-TE backbone.

Further flexibility is introduced through the combination of modes (e.g. **mode 2** + **mode 0** or **mode 2** + **mode 1**; see examples in **Supplementary Methods**) uniquely enable hybrid simulation approaches that leveraging the TE compositional realism of real genomes (**mode 2**) while expanding diversity through random synthetic non-TE backbones (**mode 0**) and introducing controllable novel TE activity to a real genome (**mode 1**), mimicking TE bursts in horizontal transfer or TE reactivation. Besides, combining TEgenomeSimulator’s digital replicas with PrinTE’s forward evolution simulation facilitates investigation in complex evolutionary history [22].

The customizable mutagenesis parameters and compositional realism of TEgenomeSimulator make it ideal for systematically evaluating tool sensitivity across identity and integrity gradients, as demonstrated in our RepeatMasker studies. Hence, TEgenomeSimulator could serve as a reference framework for benchmarking studies, establishing standardized simulation conditions for reproducible TE tool comparisons.

Future development could integrate the positional bias of TE insertion, TE dynamics over time, and epigenetic influences, bridging the gap between structural simulation and evolutionary modeling.

### Key points

- TEgenomeSimulator addresses the critical gap in large-scale genomic analysis by providing traceable, simulated genomes that serve as essential “ground-truth” benchmarks for evaluating the accuracy of TE annotation tools, particularly for non-model organisms where curated data is scarce.
- TEgenomeSimulator offers a unique methodological structure through three interoperable modes that span a spectrum of complexity: from entirely synthetic genomes (**mode 0**) and applying TEs to custom genome backbones (**mode 1**) to the high-fidelity approximation of empirical TE landscapes derived from source genomes (**mode 2**).
- TEgenomeSimulator allows for versatile parameterization of TE mutagenesis, including sequence identity, integrity, and copy number. The tool captures greater evolutionary complexity and genomic realism than existing simulators such as denovoTE-eval and GARLIC.
- While this manuscript demonstrates TEgenomeSimulator’s utility via RepeatMasker, the framework is built to evaluate a wide range of TE detection and annotation methods, uncovering how specific gradients of sequence conservation and structural integrity impact performance across different algorithms.
- TEgenomeSimulator’s ability to combine simulation modes provides a tunable continuum between evolutionary abstraction and genomic realism. This flexibility, paired with its open-source design and compatibility as a “burn-in” input for forward-evolution simulators like PrinTE, ensures that it is a comprehensive resource for the genomic research community.

## Acknowledgements

We want to thank Drs Chen Wu, Julie Blommaert, Jason Shiller, and David Chagné for reviewing this manuscript and providing insightful comments.

## Author Contributions

Ting-Hsuan Chen (Conceptualization [lead], Software [lead], Analysis [lead], Writing — original draft [lead]), Olivia Angelin-Bonnet (Software [supporting], Writing — review & editing [supporting], James Bristow (Software [supporting], Writing — Review & editing [supporting]), Christopher Benson (Writing — Review & editing [supporting]), Shujun Ou (Writing — review & editing [supporting]), Cecilia Deng (Conceptualization [supporting], Writing — review & editing [supporting], Supervision [supporting]), Susan Thomson (Conceptualization [supporting], Funding acquisition [lead], Supervision [lead], Writing — review & editing [lead])

## Use of AI in this study

ChatGPT o4 and 5 were used to help draft and improve scripts during the development of TEgenomeSimulator, and to suggest edits for clarity and grammar in the manuscript.

## Data availability

TEgenomeSimulator is available at https://github.com/Plant-Food-Research-Open/TEgenomeSimulator under the GPL-3.0 license, and all codes for the analyses in this paper are available at https://github.com/Plant-Food-Research-Open/TEgenomeSimulator_publication_code under the MIT license.

