## Supplementary Figures S1-S8 for "TEgenomeSimulator: A Flexible Framework for Simulating Genomes with Configurable Transposable Element Landscapes"

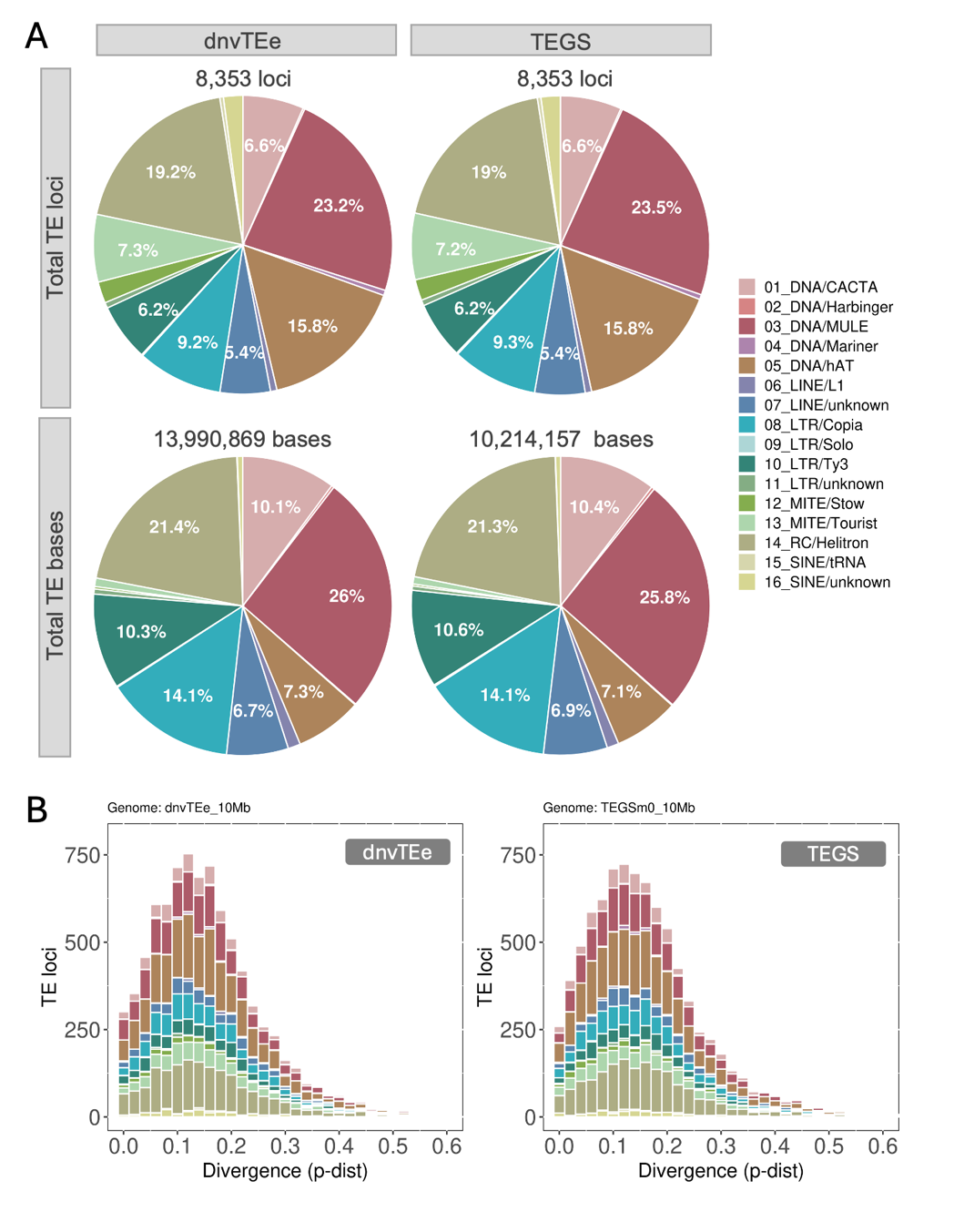

**Fig. S1. Comparison between TEgenomeSimulator and denovoTE-eval.** (A) TE composition of synthetic genomes generated by denovoTE-eval (dnvTEe) and TEgenomeSimulator (TEGS) using 10 Mb randomly simulated backbones. The dnvTEe simulation was configured using the TE mutagenesis table (‘TElib_sim_list.table’) produced by TEGS to ensure comparability. TE superfamilies representing more than 5% of the total composition are labelled. (B) Sequence divergence distributions of simulated TEs, coloured by TE superfamily.

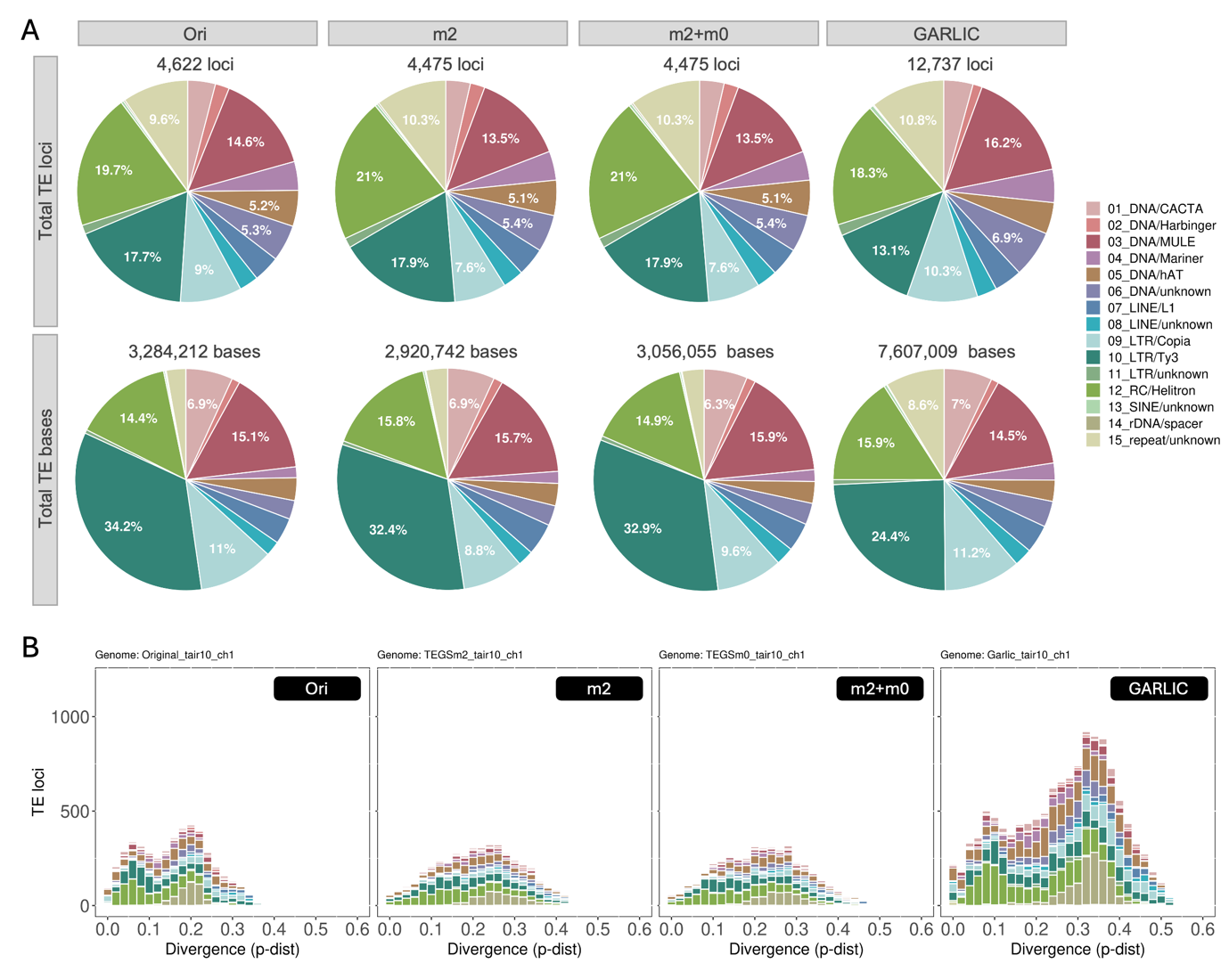

**Fig. S2. Comparison between TEgenomeSimulator and GARLIC.** (A) TE composition of the original TAIR10 chromosome 1 (Ori) and the synthetic chromosomes generated by TEgenomeSimulator mode 2 (m2), mode 2 followed by mode 0 (m2+m0), and GARLIC. TE superfamilies representing more than 5% of the total composition are labelled. (B) Sequence divergence distributions of simulated TEs, coloured by TE superfamily.

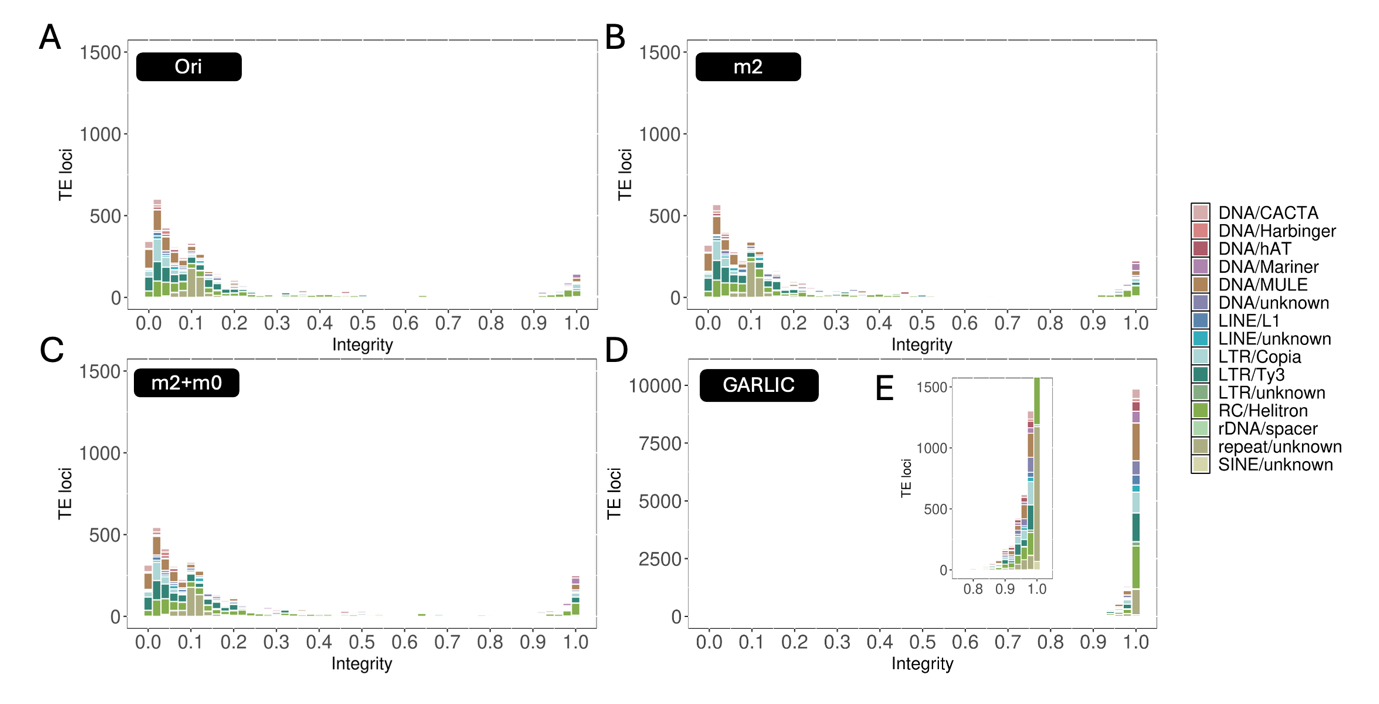

**Fig. S3. TE sequence integrity profiles.** (A-D) Distributions of TE sequence integrity in the original TAIR10 chromosome 1 (A) and,he synthetic chromosomes generated by TEgenomeSimulator mode 2 (B), mode 2 followed by mode 0 (C), and GARLIC (D). (E) Enlarged view of panel (D) highlighting the integrity range between 0.8 and 1.

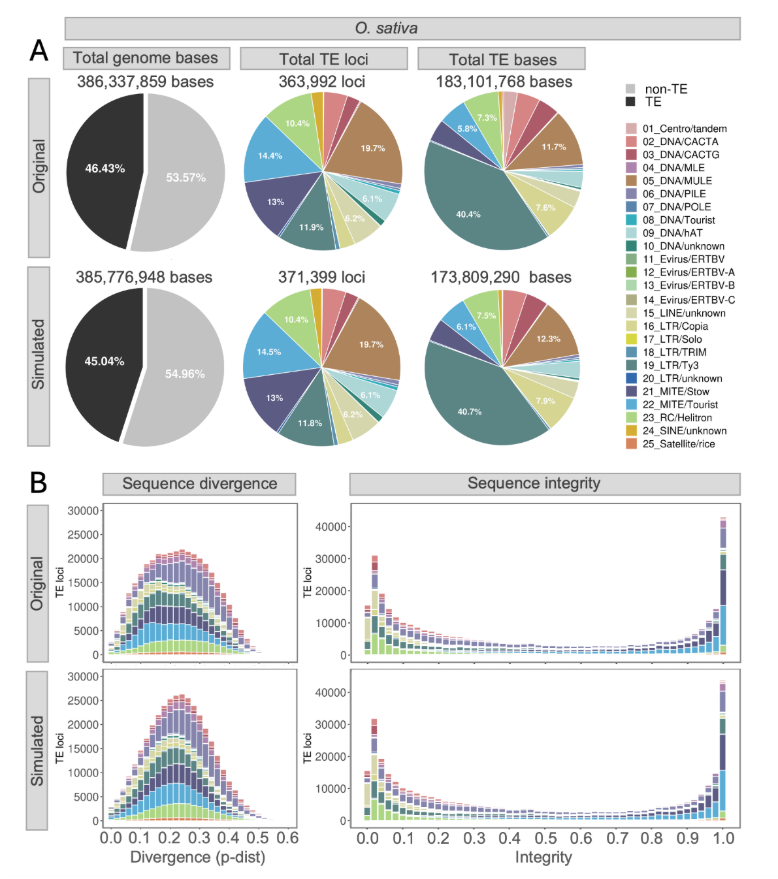

**Fig. S4**. **TE Composition Approximation for *Oryza sativa* genome.** (A) TE/non-TE sequence-occupied proportions and TE compositions (loci and bases) in the original and simulated genomes. (B) TE sequence diversity and integrity distribution colored by TE superfamily.

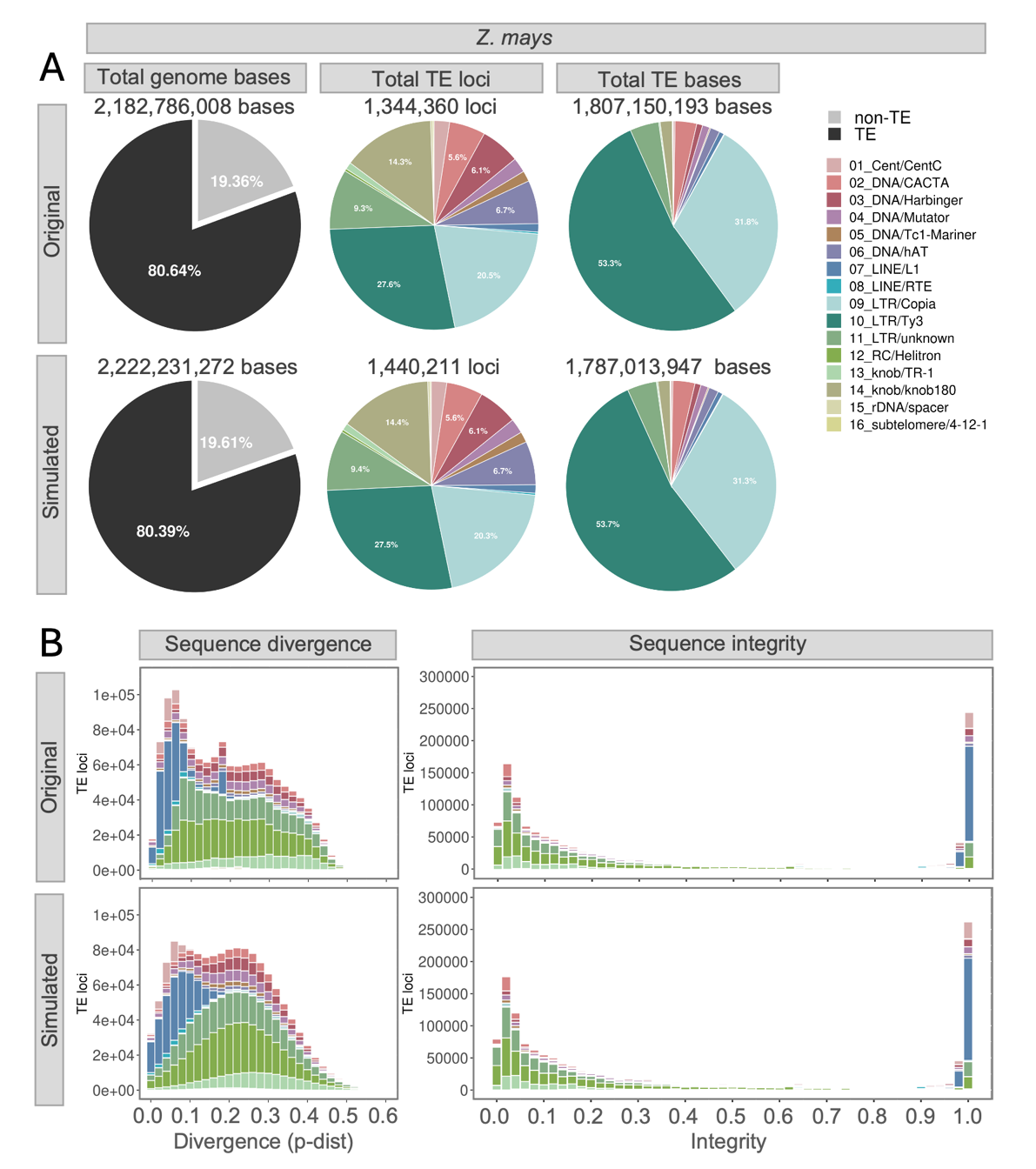

**Fig. S5**. **TE Composition Approximation for *Zea mays* genome.** (A) TE/non-TE sequence-occupied proportions and TE compositions (loci and bases) in the original and simulated genomes. (B) TE sequence diversity and integrity distribution colored by TE superfamily.

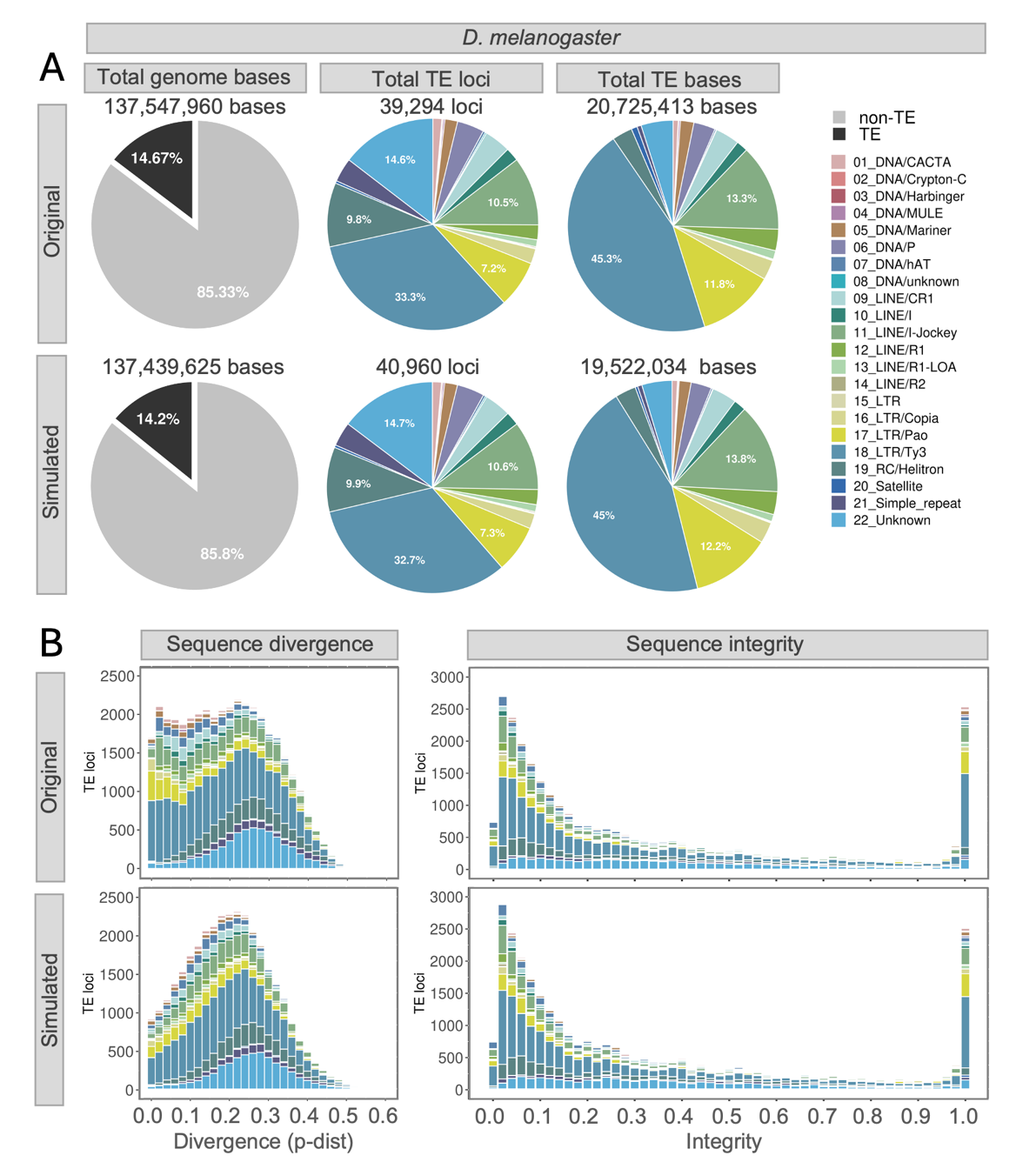

**Fig. S6**. **TE Composition Approximation for *Drosophila melanogaster* genome.** (A) TE/non-TE sequence-occupied proportions and TE compositions (loci and bases) in the original and simulated genomes. (B) TE sequence diversity and integrity distribution colored by TE superfamily.

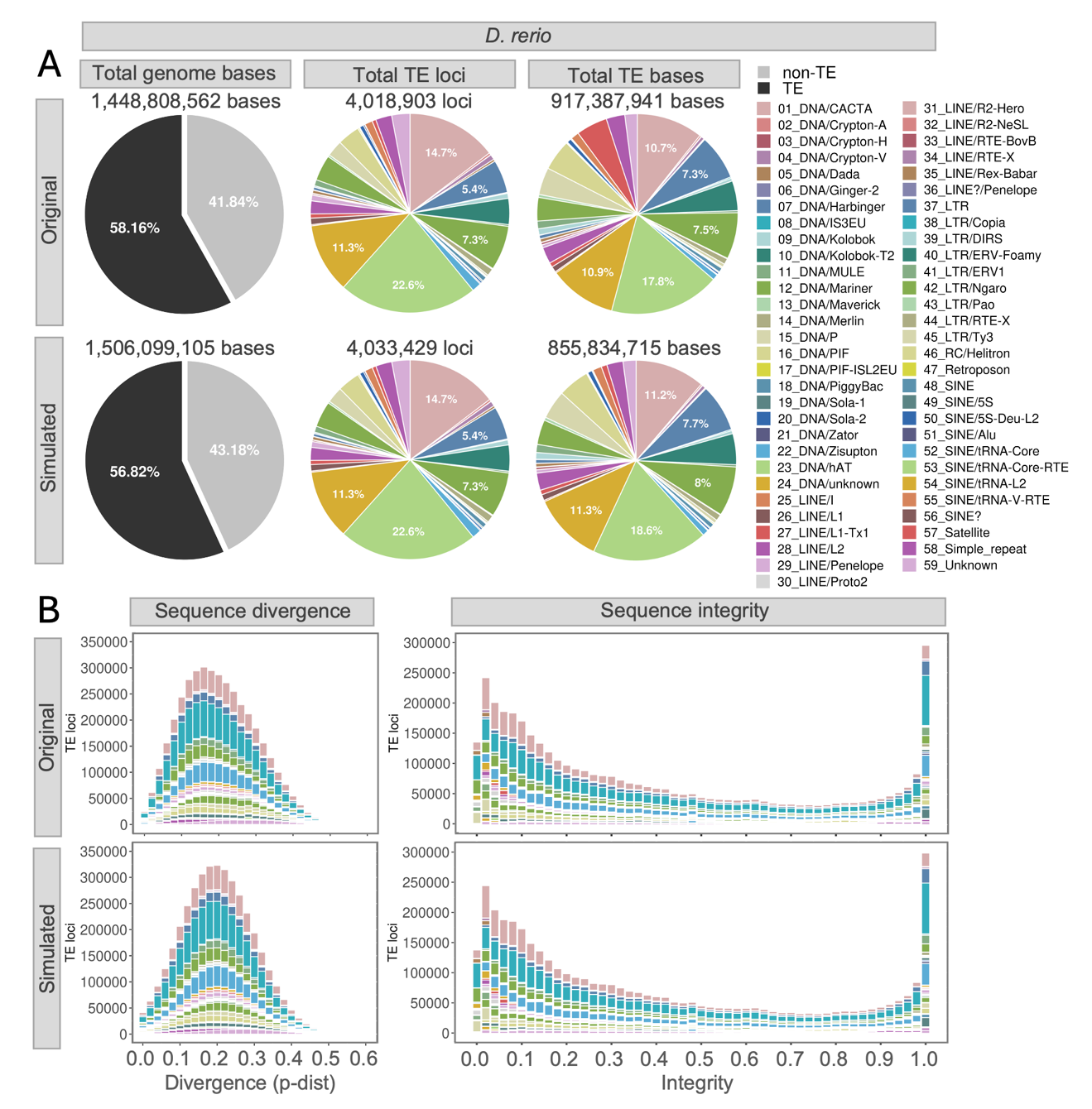

**Fig. S7**. **TE Composition Approximation for *Danio rerio* genome.** (A) TE/non-TE sequence-occupied proportions and TE compositions (loci and bases) in the original and simulated genomes. (B) TE sequence diversity and integrity distribution colored by TE superfamily.

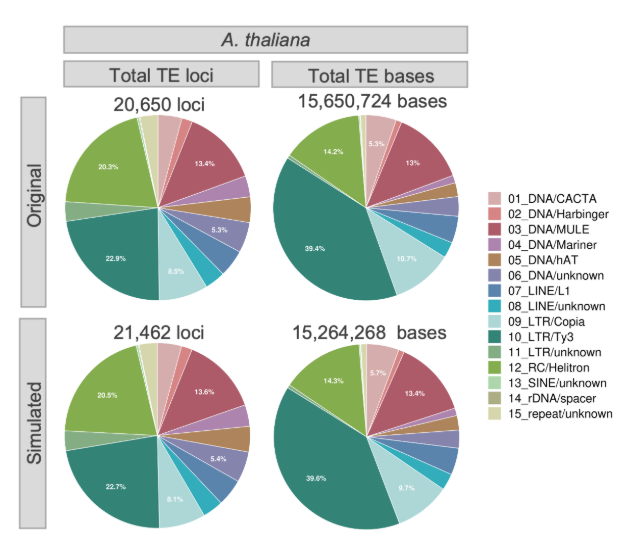

**Fig. S8**. **TE composition of the original and simulated Arabidopsis genomes.** TE superfamily proportions higher than 5% were indicated.
