## Supplementary Methods for "TEgenomeSimulator: A Flexible Framework for Simulating Genomes with Configurable Transposable Element Landscapes"

1. **Implementation**

***1.1 The Three Simulation Modes***

The Random Synthesized Genome mode (--mode 0) will synthesize chromosome sequences using a set of user-defined parameters, including chromosome length and GC content. A TE library can be provided using the ‘--repeat’ flag, which is used to simulate a population of TE sequences through random mutation, including nucleotide substitutions, 1-base InDel, target site duplication, and fragmentation. Simulated TE sequences are then randomly inserted into the synthesized chromosome sequences.

The Custom Genome mode (--mode 1) requires users to provide a ‘backbone’ genome in which TE sequences have been removed. For convenience, this mode provides an option for users to generate such a genome using the ‘--to_mask’ flag. RepeatMasker ^48^ is used to mask and remove repeats from the genome based on a TE library supplied by users. TE sequence mutation and random nested insertions are introduced in the same way as the first mode. Users must also specify a second TE library with ‘--repeat2’, which is used for TE masking and removal. Note that the simulator allows users to provide two different TE libraries for masking repeats and for TE sequence mutation, respectively.

The TE Composition Approximation mode (--mode 2) utilizes the ‘--to_mask’ flag implemented in the Custom Genome Mode, with the option of using two different TE libraries. By default, RepeatMasker output is utilized in the configuration of downstream TE mutations. In brief, the simulated TE composition is modeled on that of the original genome in terms of nucleotide substitution rate, InDel rate, integrity, and the absolute and relative abundance of TE families. The TE insertions and nested insertions in this mode are independent of the TE distribution in the original genome.

***1.2 Simulating TE Mutagenesis***

TE libraries provided by the user must adhere to the RepeatMasker-recommended format (“>repeat_family_name#class/subclass”), e.g, “>ATCOPIA10#LTR/Copia”. It is recommended that each TE family be represented by its full-length sequence in the library file. For LTR retrotransposon, this means that the LTR and internal domain should be concatenated into a single sequence in the form LTR-INT-LTR, rather than stored as separated entries. If split LTR retrotransposon entries are presented, TEgenomeSimulator automatically detects and assembles them into full-length sequences. In such cases, split entries belonging to the same LTR retrotransposon family mush share the same family name followed by the suffix “*INT” or “*LTR” immediately before the the hashtag (“#”), where “*” may be an underscore (“_”), hyphen (“-”), or omitted.

TEgenomeSimulator extracts classification information from the TE library for incorporation into a final GFF file. Metadata, including TE family name, superfamily, subclass, and sequence length, are also extracted from the TE library headers and compiled into the output file ‘TElib_sim_list.table’. This table also defines the mutagenesis parameters for each TE family, including TE family copy number, mean sequence identity (*I*), standard deviation of mean identity, and the proportion of sequence divergence (*D = 100* - *I)* contributed by InDel (*i*). The remaining proportion of divergence is attributed to nucleotide substitution (*s*), such that *i + s = 100*. The table also includes values for target site duplication (TSD) length and proportions of fragmented TE loci and nested TE insertions, each relative to the total TE copy number. The assignment of copy number, sequence identity, and sequence integrity are further described below.

***1.2.1 Copy number assignment.*** In both Random Synthesized Genome and Custom Genome modes, the user is required to provide a value range for the random assignment of a copy number to each TE family. The TE Composition Approximation mode, however, uses the copy number information of each family obtained from RepeatMasker as the exact copy number value.

***1.2.2 Sequence identity****.* Both the Random Synthesised and Custom Genome modes assign sample sequence identity values from predefined normal distributions. The mean value and the standard deviation of the mean are, by default, randomly selected from ranges between 80 to 95 (%) and 1 to 20 (%), respectively. For the Genome Approximation mode, the mean and standard deviation of sequence identity distributions are derived from the RepeatMasker output for each TE family member. For each TE insertion, an integer is then randomly drawn from the generated distribution, regarded as the original identity of the family sequence. This value is further adjusted, following the approach implemented in denovoTE-eval [37], by adding 50% of the difference between the sampled identity value and 100. It is calculated as *I' = I + (100 -* *I) × 0.5,* where *I* is the original identity sampled from the normal distribution, and *I'* is the adjusted identity value. This adjustment causes a skew of the identity distribution to the right (i.e., closer to 100%), reflecting that TE sequences with high divergence from the family sequence are less likely to be recognized as family members.

***1.2.3 Target Site Duplication (TSD)****.* TSD length is implemented based on literature findings, while the sequence of TSD is randomly synthesized. The simulator then randomly assigns the TE member to forward or reverse strands before performing sequence fragmentation and integration.

***1.2.4 Sequence integrity****.* Both Random Synthesized Genome and Custom Genome modes implement a beta distribution to model the integrity of TE sequences. The default alpha and beta values are 0.5 and 0.7, respectively, to create an asymmetric U-shape distribution in which the probability of low sequence integrity (e.g., <10%) is higher than that of high integrity (e.g.,>90%). Alpha and beta values can be adjusted using the --alpha (or -a) and --beta (or -b) parameters. The integrity value for a TE member is sampled from the established beta distribution and then converted to a number representing the bases to be removed from the 5’ end of that sequence. For example, if a TE member is 1 kbp at its full length and is assigned with 70% integrity, the simulator would remove 300 bases, which accounts for 30% of the intact length, from its 5’ end**.** By default, only 0.1% of TE copies in each TE family remain untouched, ensuring the representation of intact TEs in the simulation. For sequence integrity in the TE Composition Approximation mode, TEgenomeSimulator utilizes a randomly sampled value from a sequence integrity distribution extracted from the RepeatMasker output.

***1.3 Random and Nested TE Insertion***

TEgenomeSimulator randomly samples a value between 0 and 30% as the default proportion of TE copy number to form nested insertions. This is only applied to LTR retrotransposons and is implemented across all simulation modes. The simulator performs simple TE insertion before executing nested TE insertion. Target sites for simple insertion and target TEs for nested insertion are randomly chosen.

***1.4 Output Files of TEgenomeSimulator***

TEgenomeSimulator generates a series of output files that are useful for downstream analysis, including the fasta file of the simulated genome, the fasta file of all TE sequences inserted in the simulated genome, and the TE annotation file in GFF format that records the sequence location, identity, integrity, and information to link the nested insertion to disrupted TEs.

2. **Use-case examples**

***2.1 Random sequence simulation differing in size of non-TE backbone (mode 0)***

```
#!/bin/bash
echo "chr1,10000,35" > random_genome_chr_index_10k.csv
echo "chr1,100000,35" > random_genome_chr_index_100k.csv
echo "chr1,1000000,35" > random_genome_chr_index_1000k.csv
echo "chr1,10000000,35" > random_genome_chr_index_10000k.csv
echo "chr1,100000000,35" > random_genome_chr_index_100000k.csv
echo "chr1,100000000,35" > random_genome_chr_index_1000000k.csv
echo "chr2,100000000,35" >> random_genome_chr_index_1000000k.csv
echo "chr3,100000000,35" >> random_genome_chr_index_1000000k.csv
echo "chr4,100000000,35" >> random_genome_chr_index_1000000k.csv
echo "chr5,100000000,35" >> random_genome_chr_index_1000000k.csv
echo "chr6,100000000,35" >> random_genome_chr_index_1000000k.csv
echo "chr7,100000000,35" >> random_genome_chr_index_1000000k.csv
echo "chr8,100000000,35" >> random_genome_chr_index_1000000k.csv
echo "chr9,100000000,35" >> random_genome_chr_index_1000000k.csv
echo "chr10,100000000,35" >> random_genome_chr_index_1000000k.csv

repeat=your_TE_lib.fasta
min=1
max=10

for chr in 10 100 1000 10000 100000 1000000
do
indcsv=random_genome_chr_index_${chr}k.csv
prefix=m0_chr${chr}k
cd $OUT
apptainer run TEgenomeSimulator_v1.0.0.sif -M 0 -p $prefix -g $genome -r $repeat -o ".”
done
```

***2.2 Impact of TE copy number on simulated genome size (mode 1)***

```
#!/bin/bash
genome=your_real_genome_nonTE.fa
repeat=your_curated_TElib.fa

MIN=(1 5 5 5 5)
MAX=(10 100 500 1000 2000)
for((i=0;i<${#MIN@]};i++))
do
min=${MIN[$i]}
max=${MAX[$i]}
prefix=m1_cn_${min}_${max}
apptainer run TEgenomeSimulator_v1.0.0.sif -M 1 -p $prefix -g $genome -r $repeat -m $max -n $min -o ".”
done
```

***2.3 Impact of alpha and beta values (mode 1)***

```
#!/bin/bash
A=(0.5 0.5 0.5 0.75 1)
B=(0.5 0.75 1 0.5 0.5)
min=5
max=100
genome=your_real_genome_nonTE.fa
repeat=your_curated_TElib.fa
for((i=0;i<${#A@]};i++))
do
alpha=${A[$i]}
beta=${B[$i]}
prefix=m1_itg_${i}
apptainer run TEgenomeSimulator_v1.0.0.sif -M 1 -p $prefix -g $genome -r $repeat -m $max -n $min -a $alpha -b $beta -o ".”
done
```

***2.4 Impact of mean sequence identity (mode 1)***

```
#!/bin/bash
MINIDN=(90 80 70 70 70 70 70)
MAXIDN=(100 90 80 85 90 95 100)
min=5
max=100
genome=your_real_genome_nonTE.fa
repeat=your_curated_TElib.fa
for((i=0;i<${#MINIDN@]};i++))
do
minidn=${MINIDN[$i]}
maxidn=${MAXIDN[$i]}
prefix=m1_idn_${minidn}_${maxidn}
apptainer run TEgenomeSimulator_v1.0.0.sif -M 1 -p $prefix -g $genome -r $repeat -m $max -n $min --maxidn $maxidn --minidn $minidn -o ".”
done
```

***2.5 Combinational impact of sequence identity and standard deviation (mode 1)***

```
#!/bin/bash
MINIDN=(90 70 70 90)
MAXIDN=(100 80 80 100)
MINSD=(15 15 1 1)
MAXSD=(20 20 5 5)

min=5
max=100
genome=your_real_genome_nonTE.fa
repeat=your_curated_TElib.fa
for((i=0;i<${#MINIDN@]};i++))
do
minidn=${MINIDN[$i]}
maxidn=${MAXIDN[$i]}
minsd=${MINSD[$i]}
maxsd=${MAXSD[$i]}

prefix=m1_scenario_${i}
apptainer run TEgenomeSimulator_v1.0.0.sif -M 1 -p $prefix -g $genome -r $repeat -m $max -n $min --maxidn $maxidn --minidn $minidn –maxsd $maxsd --minsd $minsd -o ".”
done
```

***2.6 A digital replica of A. thaliana (mode 2)***

```
#!/bin/bash
genome=GCF_000001735.4_TAIR10.1_genomic.fna
repeat=athrep.updated.nonredun.noCenSatelli.fasta
threads=8 #for RepeatMasker
prefix=tair10
apptainer run TEgenomeSimulator_v1.0.0.sif -M 2 -p $prefix -g $genome -r $repeat -r2 $repeat -o ".” -t $threads
```

***2.7 TE replica of A thaliana chromosome 1 in random sequence (mode 2 + 0)***

```
#!/bin/bash
# Run mode 2 on TAIR10 chr1
genome=GCF_000001735.4_TAIR10.1_genomic_chr1.fna
repeat=athrep.updated.nonredun.noCenSatelli.fasta
threads=8 #for RepeatMasker
prefix=tair10_chr1_m2
apptainer run TEgenomeSimulator_v1.0.0.sif -M 2 -p $prefix -g $genome -r $repeat -r2 $repeat -o ".” -t $threads

# Extract size (use samtools faidx) and GC% (use emboss’ tool: infoseq) of the non-TE chr1
m2_result_dir=./TEgenomeSimulator_tair10_chr1_m2_result
TEGS_processed_nonTE_chr=$m2_result_dir/ GCF_000001735.4_TAIR10.1_genomic_chr1.fna.masked.reformatted.nonTE.emptfixed
samtools faidx $TEGS_processed_nonTE_chr
nontesize=$(awk '{print $2}' GCF_000001735.4_TAIR10.1_genomic_chr1.fna.masked.reformatted.nonTE.emptfixed.fai
gc=$(cat $TEGS_processed_nonTE_chr | infoseq -auto -only -name -length -pgc stdin | awk 'NR>1{print $3}')
echo "chr1,$nontesize,$gc" > tair10_chr1_index_nonte.csv

# Run mode 0 using chr information (size and GC%), TE table,
# and processed TE library from mode 2’s outputs
indcsv=tair10_chr1_index_nonte.csv
repeat=$m2_result_dir/athrep.updated.nonredun.noCenSatelli.fasta.stitched
tetable=$m2_result_dir /TElib_sim_list_mode2.table
prefix=tair10_chr1_m0
apptainer run TEgenomeSimulator_v1.0.0.sif -M 0 -p $prefix -c $indcsv -r $repeat –te_table $tetable -frag_mode 2 -o ".”
```

***2.8 Adding recent Copia burst to digital replica of A. thaliana chr1 (mode 2 + 1)***

```
#!/bin/bash
# Run mode 2 on TAIR10 chr1
genome=GCF_000001735.4_TAIR10.1_genomic_chr1.fna
repeat=athrep.updated.nonredun.noCenSatelli.fasta
threads=8 #for RepeatMasker
prefix=tair10_chr1_m2
apptainer run TEgenomeSimulator_v1.0.0.sif -M 2 -p $prefix -g $genome -r $repeat -r2 $repeat -o ".” -t $threads

# Run mode 1 using pre-processed TE library that
# only contains the active TE families
genome= $m2_result_dir/ GCF_000001735.4_TAIR10.1_genomic_chr1.fna.masked.reformatted.nonTE.emptfixed
repeat=athrep.updated.nonredun.noCenSatelli.Copia.fasta
prefix=tair10_chr1_m1_copia_burst
minidn=90
maxidn=100
minsd=1
maxsd=5
apptainer run TEgenomeSimulator_v1.0.0.sif -M 1 -p $prefix -g $genome -r $repeat -m $max -n $min --maxidn $maxidn --minidn $minidn –maxsd $maxsd --minsd $minsd -o ".”
```
